# Age-associated Autoimmunity Driven by T cell Immunosenescence

**DOI:** 10.64898/2026.01.07.698250

**Authors:** Ethan C. McCarthy, Maryamsadat Seyedsadr, Madison F. Bang, Caitlyn L. H. Pham, Christian Bustillos, Isabela Salazar, Ivan A. Salladay-Perez, Jason K. Whitmire, Christopher Fu, Sophie Blanc, Mehdi Bouhaddou, Jocelyn Weng, Faranak Fattahi, Melissa G. Lechner, Perry Shieh, Anthony J. Covarrubias, Maureen A. Su

## Abstract

While certain autoimmune conditions are more common in the young (e.g. Type 1 Diabetes), others are more frequent in the aged. A striking example is chronic inflammatory demyelinating polyneuropathy (CIDP), a CD4+ T cell-mediated autoimmune disease of the peripheral nervous system, which has a peak decade of onset of 70-79 years. How aging predisposes to autoimmunity, however, remains unclear. CD4+ T cells are highly susceptible to age-associated changes, including acquisition of immunosenescent features such as enhanced SA-β-gal activity, increased Cdkn1a (p21) expression, and upregulation of Tnfsf8 (CD153). We show here that CD4+ T cells exhibiting these changes are increased in CIDP patients and mice with CIDP-like disease. These CD4+ T cells exhibit a senescence-associated secretory phenotype (SASP), show functional senescence (i.e., decreased proliferation and resistance to apoptosis), and have enhanced capacity for inciting neuropathy. Notably, a senescent cell-clearing senolytic agent (fisetin) decreased the pathogenic capacity of CD4+ T cells, and a SASP-suppressing senomorphic therapy (ruxolitinib) protected mice against autoimmune demyelination. Together, these findings delineate a key role for age-associated senescent CD4+ T cells in driving age-associated autoimmunity.

## Introduction

Increasing age is a risk factor for the development of many autoimmune disorders^1^ ^2^, including autoimmune peripheral neuropathies. For instance, CIDP is a debilitating autoimmune demyelinating disease of the peripheral nervous system (PNS)^3, 4^ that most often manifests in the eighth decade of life^3^. Patients with CIDP experience progressive weakness, sensory dysfunction, and loss of mobility, with severe cases leading to profound long-term disability and, rarely, death. With existing therapies, few patients achieve stable disease off treatment and more than half require ongoing immunosuppression^4^. Similarly, Guillain-Barré Syndrome (GBS), often considered the acute counterpart of CIDP, occurs more frequently in older individuals and shows a 20% increase in incidence with every 10 year increase in age^5^. The mechanisms underlying this age-associated autoimmunity risk remain poorly understood, and delineating these pathways could provide potential therapeutic targets to stop disease onset or progression.

CD4+ T cells play a key role in the autoimmune destruction of the PNS and are prominent within the infiltrated peripheral nerves of CIDP patients and mice with CIDP-like disease^6, 7, 8, 9^. Transfer of CD4+ T cells derived from CIDP mice are sufficient to incite neuropathy in immunodeficient hosts^7, 8^, highlighting the importance of CD4+ T cells in CIDP pathogenesis. Intriguingly, recent evidence suggests that CD4+ T cells are highly susceptible to age-associated changes, a potential consequence of thymic involution, chronic antigen stimulation, and other cellular stressors^10, 11^. For instance, aging is associated with contraction of the naïve T cell compartment with concomitant expansion of effector/memory T cell proportions^12^. With age, T cells also acquire cellular senescence features, including increased β-galactosidase (β-gal) activity, proliferative arrest, resistance to apoptosis, and secretion of pro-inflammatory factors resembling a senescence-associated secretory phenotype (SASP)^11, 12, 13^. In addition, age-associated CD4+ T cells upregulate cell type-specific age-associated markers, such as the signaling molecule Tnfsf8 (CD153)^14, 15, 16, 17^. Thus, CD4+ T cells are a key pathogenic cell type in PNS autoimmunity that undergo profound changes with aging.

Notably, recent evidence has demonstrated that these age-associated changes in CD4+ T cells are functionally consequential, with both beneficial and deleterious outcomes. For example, age-associated alterations in CD4+ T cells have been reported to enhance anti-cancer immunity to facilitate longevity and healthy aging^18^. At the same time, age-dependent changes in CD4+ T cells may also have detrimental effects such as promoting the formation of tertiary lymphoid structures to exacerbate kidney disease^14^. Thus, age-associated changes in CD4+ T cells have diverse and profound consequences. However, it remains unclear the impact of age-associated changes in CD4+ T cells on the development of age-associated PNS autoimmunity.

Here, we demonstrate that CD4+ T cells exhibiting age-associated senescent features accumulate in CIDP patients and in a CIDP mouse model. These CD4+ T cells are highly pathogenic, demonstrating an increased capacity to transfer neuropathy. Mechanistically, these CD4+ exhibit a pro-inflammatory SASP, which spreads senescence to nearby peripheral nerve Schwann cells. Depletion of senescent CD4+ T cells with a senolytic agent (fisetin) impairs their autoimmune capacity, and targeting the CD4+ T cell SASP in CIDP mice with a senomorphic agent (ruxolitinib) protects against autoimmune peripheral neuropathy. Together these findings point to a key role for age-associated CD4+ T cell senescence in driving CIDP pathogenesis and suggest the deployment of serotherapies as a class of medications in preventing aging-associated autoimmune diseases.

## Results

### CD4+ T cells in CIDP patients exhibit immunosenescence features

Acquisition of age-associated dysfunction, exhaustion, and senescence features in immune cells, termed immunosenescence, has been linked to age-associated neurodegenerative disorders (e.g. Alzheimer’s Disease and Parkinson’s Disease)^19^, but its role in age-associated autoimmune neuropathies has not yet been studied. Given that we and others previously established a critical role of CD4+ T cells in the pathogenesis of CIDP^7, 8, 20^, we first assessed a key biomarker of cellular senescence, senescence associated beta-galactoside (SA-β-Gal) activity^11, 12^, in CD4+ T cells from CIDP patients. Supporting the possibility that CIDP patients accumulate senescent CD4+ T cells, peripheral blood CD4+ T cells from CIDP patients showed increased SA-β-Gal activity relative to age-matched, healthy controls (**Figure 1A)**.

**Figure 1:**
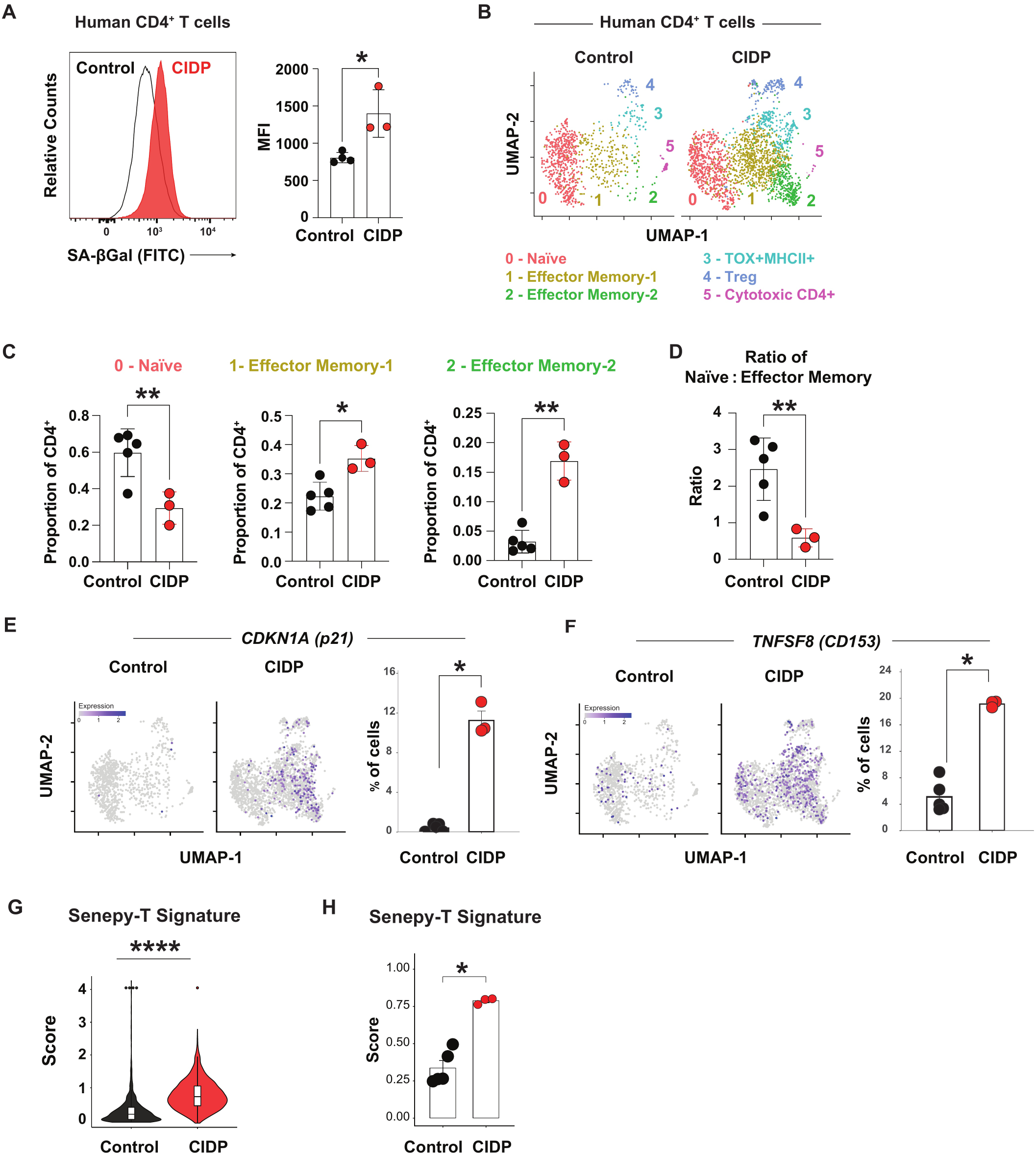
CD4+ T cells from patients with CIDP exhibit features of immunosenescence. **A)** Representative and mean fluorescence intensity (MFI) quantification of senescence-associated beta-galactosidase activity in viable, peripheral human CD3+CD4+ T cells (n=3 CIDP, n=4 age-matched healthy controls [Control]) (* P < 0.05 by two-tailed unpaired t test). Each data point represents one patient. **B)** Uniform manifold approximation and projection (UMAP) of scRNA-seq data from CIDP and age-matched healthy control peripheral CD4 T cells. **C)** Comparison of cluster proportions between groups (*P< 0.05, ** P < 0.01 by two-tailed unpaired t test with Welch’s correction). **D)** Ratio of naive (cluster 0) to effector (clusters 1 and 2) proportions (** P < 0.01 by two-tailed unpaired t test with Welch’s correction). **E,F)** Ordered feature plots and mean % frequency of CDKN1A (p21) (E) or TNFSF8 (CD153) (F) expression, split by patient group. Each point in graph represents individual patient (Wilcoxon *p<0.05). **G)** SenePy senescence scores for human blood T cell signature in CD4+ cells from healthy controls and CIDP patients. Each dot represents a cell. Boxes showed median and interquartile range (*** P< 0.0001, Mann-Whitney). **H)** SenePy senescence scores for human blood T cell signature in CD4+ cells from healthy controls and CIDP patients. Each dot represents one donor sample (* P< 0.05, Mann-Whitney, mean ±SEM).

To further delineate CD4+ T cell alterations in CIDP, we performed scRNA-seq (single cell RNA sequencing) on peripheral blood CD4+ T cells from CIDP patients (n=3, 2062 cells in total) and integrated these data with an existing, age-matched healthy control scRNA-seq dataset (n=5, 915 cells in total)^21^. Clustering analysis identified six CD4+ T cell subsets (**Figure 1B, Supplementary Table 1**) including naïve (cluster 0; **Figure S1A**: *SELL* [CD62L]^+^, *CCR7*^+^, *CD27*^+^, *TCF7* [TCF1]^+^, *CD44*^-^); effector memory (cluster 1, **Figure S1B**: *TNFRSF4* [OX40]^+^, *PRDM1* [BLIMP1]^+^, *CD40LG*^+^, *CXCR5*^+^; cluster 2, **Figure S1C:** eomesodermin [*EOMES*], T-bet [*TBX21*], Granzyme K [*GZMK*]; and interferon *[IFN]-*γ *AS1*, *CXCR3*, *CXCR6*). Consistent with known age-associated changes^11, 12^, we observed contraction of the naïve T cell subset and expansion of the effector/memory T cell subsets in CIDP individuals relative to healthy controls (**Figure 1C**), with a decreased ratio of naïve to effector CD4+ T cells in CIDP (**Figure 1D**). Moreover, expression of the canonical senescence-associated cell cycle inhibitor *CDKN1A (*p21)^13^ (**Figure 1E)** and senescence-associated signaling molecule *TNFSF8* (CD153)^16^ were higher in CIDP CD4+ T cells (**Figure 1F)**. Thus, scRNAseq analyses show alterations in CD4+ T cell subsets and changes in senescence-associated marker expression suggestive of increased CD4+ T cell senescence in CIDP.

Senescence gene signatures have proven invaluable for identifying cells with age-associated changes^22, 23, 24^. A ‘SenePy-T’ gene signature, for instance, is available which characterizes *in vivo* senescence in human peripheral blood T cells^24^. Utilizing this approach, we found significant enrichment of the ‘SenePy-T’ gene signature in CD4+ T cells from CIDP patients, compared to healthy controls (**Figure 1G**). In parallel, ‘pseudobulk’ analysis also revealed upregulation of ‘SenePy-T’ signature in CIDP (**Figure 1H**). Taken together, multiple lines of evidence demonstrate that peripheral blood CD4+ T cells from CIDP patients exhibit features of cellular senescence that distinguish them from age-matched, healthy controls.

### Senescence-associated CD153+ CD4+ T cells accumulate in peripheral nerves of a mouse with CIDP-like disease

Mechanistic insights into CIDP pathogenesis have been revealed through studies in mice that develop CIDP-like disease^27, 28^. To understand the role of senescent CD4+ T cells in PNS autoimmunity, we thus turned to the *NOD.Aire^GW/+^* mouse model of CIDP^8, 25, 26, 27, 28^. *NOD.Aire^GW/+^*mice harbor a dominant-negative mutation in the *Aire* (*Autoimmune Regulator*) gene that leads to development of spontaneous autoimmune peripheral neuropathy that shares multiple features with human CIDP (e.g., development without a trigger, peripheral nerve infiltration with CD4+ T cells)^8^. Of note, *NOD.Aire^GW/+^* mice are unique among inflammatory peripheral neuropathy models in having patient counterparts with causative mutations in the orthologous gene (*AIRE*)^29^. Additionally, because aging has been associated with loss of Aire function^30, 31^, we reasoned that Aire deficiency in the *NOD.Aire^GW/+^* mouse model may recapitulate this age-associated component of autoimmunity.

We first asked whether senescent T cells are enriched in peripheral nerves of neuropathic *NOD.Aire^GW/+^* mice. Analysis of scRNA-seq datasets of CD4+ T cells within the peripheral nerves of neuropathic *NOD.Aire^GW/+^*mice^32^ and healthy *NOD.WT* controls^33^ revealed five populations (**Figure 2A, Supplementary Table 2**), including a subset of CD4+ T cells (cluster 2) that differentially upregulates the aging-associated genes *Tnfsf8* (CD153) and *Pdcd1* (PD1) **(Figure 2B,C)**^14, 16, 34^. Additionally, senescent cells acquire expression of pro-inflammatory factors that are part of the senescence-associated secretory phenotype (SASP), and Cluster 2 upregulates multiple SASP factors [*Ifng, Il21, Il10,* and *Spp1* (osteopontin)] **(Figure 2B,D)**^11, 12^, further pointing to cluster 2’s identity as a senescence-associated population. Notably, proportion analysis revealed over-representation of cluster 2 in CD4+ T cells of *NOD.Aire^GW/+^* mice, compared to *NOD.WT* controls **(Figure 2E),** suggesting accumulation of this CD153+ PD1+ SASP-expressing population in nerves of neuropathic *NOD.Aire^GW/+^*mice.

**Figure 2:**
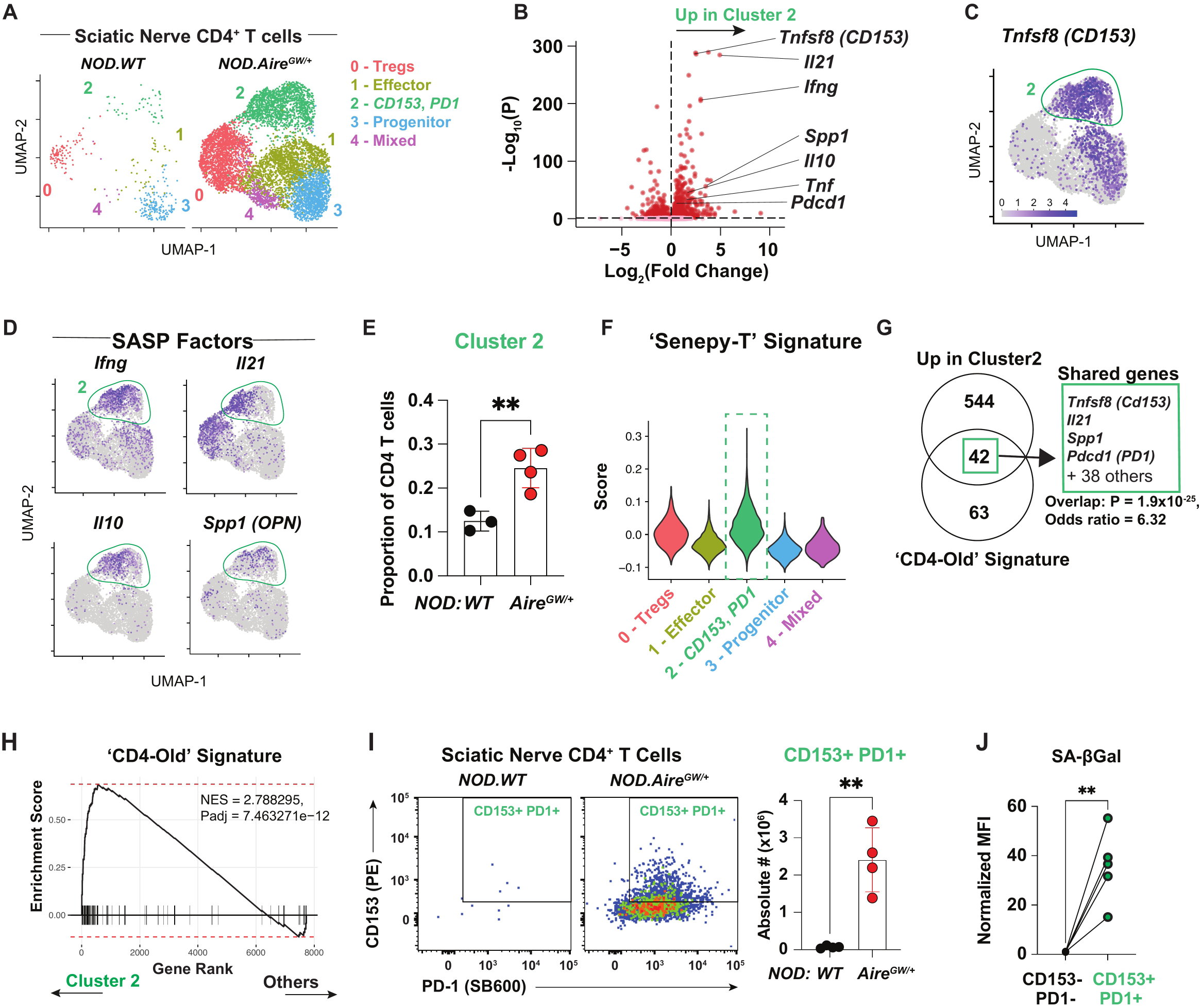
Accumulation of CD4+ T cells expressing senescent features in peripheral nerves of neuropathic mice. **A)** UMAP of scRNA-seq data from sciatic nerve-infiltrating CD4+ T cells of *NOD.WT*(n=3) and *NOD.Aire^GW/+^* (n=4) mice. Volcano plot of differentially expressed genes upregulated by cluster 2 (Log2FC < 0.5; Padjj<0.05) of the integrated sciatic nerve CD4+ T cell scRNA-seq dataset, relative to all other clusters. **C,D)** Ordered feature plot of Tnfsf8 (CD153) and SASP factor expression. Cluster 2 is outlined in green. **E)** The proportion of CD4 T cells belonging to cluster 2 for each sample within the scRNA-seq dataset, split by genotype. **F)** SenePy-T gene signature scores in clusters. **G)** Venn diagram analysis using Fisher’s test of genes in the ‘CD4-Old’ signature and genes differentially upregulated in cluster 2 (Padj<0.05; Log2FC>0.5). **G)** Venn diagram analysis using Fisher’s test of genes in the ‘CD4-Old’ signature and genes differentially upregulated in cluster 2 (Padj<0.05; Log2FC>0.5). **H)** GSEAof ‘CD4-Old” signature in genes differentially upregulated in cluster 2.**1)** Representative flow cytometry and quantification of the absolute number of CD153+PD-1 + CD4+ T cells within the sciatic nerve of *NOD.WT* and *NOD.Aire^GW/+^* mice (** P < 0.01 by two-tailed unpaired t test). Bar graph shows the absolute number from each mouse and mean ± 1 standard deviation. **J)** Normalized mean fluorescence intensity (MFI) quantification by flow cytometry of SA-pGal activity expression in *NOD.Aire^GW/^*^+^ peripheral nerve-infiltrating CD153+ PD1+and CD153-PD1- CD44+ CD4+ T cells (*P < 0.05, **P < 0.01 by paired, two-tailed t test). Pairs represent cells gated from the same mouse.

Given cluster 2’s overrepresentation in *NOD.Aire^GW/+^*mice, we further confirmed cluster 2’s identity as a senescent CD4+ T cell population using the ‘Senepy-T’ signature^24^, which was highest in cluster 2 (**Figure 2F**). Because the ‘SenePy-T’ signature contains both CD4+ and CD8+ T cell samples, however, this signature is not specific for age-associated CD4+ T cells. To address this, we established an aging-associated core signature specific for CD4+ T cells, using CD4+ T cells from a single-cell atlas of the aging mouse (Tabula Muris Senis)^34^ (**Figure S2A**). Comparison of old (18-30 months old, n=10) vs. young (1-3 months old, n=3) mice revealed differential upregulation of *Tnfsf8 (Cd153), Pdcd1 (PD1)*, and SASP factors (e.g. *Il21, Spp1*) (**Figure S2B,C**), which were used to establish a core gene signature (termed ‘CD4-Old’) for age-associated CD4+ T cells (**Supplementary Table 3, Figure S2D;** Padj<0.05; Log2FC>1). Cross-referencing of ‘CD4-Old’ signature genes (89 genes) with cluster 2 markers (570 genes) revealed a strong overlap, with 42 shared genes including *Tnfsf8 (Cd153), Pdcd1 (PD1)* and SASP factors (*Il21, Spp1*) (**Figure 2G**). Supporting cluster 2’s identity as senescent CD4+ T cells, GSEA showed significant enrichment of the ‘CD4-Old’ gene signature within cluster 2, compared to other clusters (**Figure 2H**). Taken together, these findings substantiate cluster 2’s identity as an age-associated senescent CD4+ T cell population.

Finally, we utilized flow cytometric analysis to quantify peripheral nerve-infiltrating senescence-associated CD4+ T cells. Consistent with our scRNAseq data, CD153+ PD1+ CD4+ T cells were increased in sciatic nerves of *NOD.Aire^GW/+^* mice, compared to *NOD.WT* mice (**Figure 2I**), suggesting their accumulation in *NOD.Aire^GW/+^* mice. Furthermore, these cells showed higher senescence-associated (SA)-β-gal activity **(Figure 2J)** compared to CD153-PD1- CD4+ T cell controls, supporting our transcriptomic data that CD153+ PD1+ CD4+ T cells exhibit key senescence features **(Figure 2B-D, F-H)**. Taken together, these data collectively demonstrate a higher burden of senescence-associated CD153+ PD1+ CD4+ T cells within peripheral nerves of neuropathic *NOD.Aire^GW/+^* mice.

### CD153+ CD4+ T cells exhibit a functional phenotype associated with senescence

We next sought to determine whether CD153+ PD1+ CD4+ T cells exhibit senescence-associated functional changes. Senescent cells are characterized by reduced proliferative capacity^35^, and we noted higher protein expression of senescence-associated *Cdkn1a* (p21), a key inhibitor of proliferation, in CD153+ PD1+ CD4+ T cells from sciatic nerves of *NOD.Aire^GW/+^* mice **(Figure 3A, left and center).** Thus, we sought to assess the proliferative capacity of CD153+ PD1+ CD4+ T cells. Comparison of CellTrace Violet proliferation dye staining showed lower dilution of dye in sorted CD153+ PD1+ CD4+ T cells, compared to CD153- PD1- controls **(Figure 3A, right)**, demonstrating lower proliferative capacity. In addition to reduced proliferative capacity, senescent cells are also characterized by resistance to apoptosis via upregulation of genes associated with senescent cell anti-apoptotic pathways^35^. Thus, we assessed apoptotic capacity of CD153+ PD1+ CD4+ T cells *in vitro*. After 3 days of culture, lower levels of the apoptotic marker Annexin V were seen in CD153+ PD1+ CD4+ T cells, compared to CD153- PD1- counterparts **(Figure 3B**), suggesting that CD153+ PD1+ CD4+ T cells are resistant to apoptosis. In support of these findings, bulk RNA sequencing (RNAseq; **Supplementary Table 4; Figure S3A**) of sorted CD153+ PD1+ CD4+ T cells vs. CD153- PD1+ CD4+ T cells from neuropathic *NOD.Aire^GW/+^* mice (n=3) showed upregulation of the cell cycle inhibitor *Cdkn1a* (p21) (**Figure 3C,D**) and anti-apoptotic factors, *Src* and *Bcl2ls (Bcl-xl)* (**Figure 3C,E**). Thus, CD153+ PD1+ CD4+ T cells in *NOD.Aire^GW/+^* mice exhibit anti-proliferative and anti-apoptotic functional features, consistent with a senescence phenotype.

**Figure 3.**
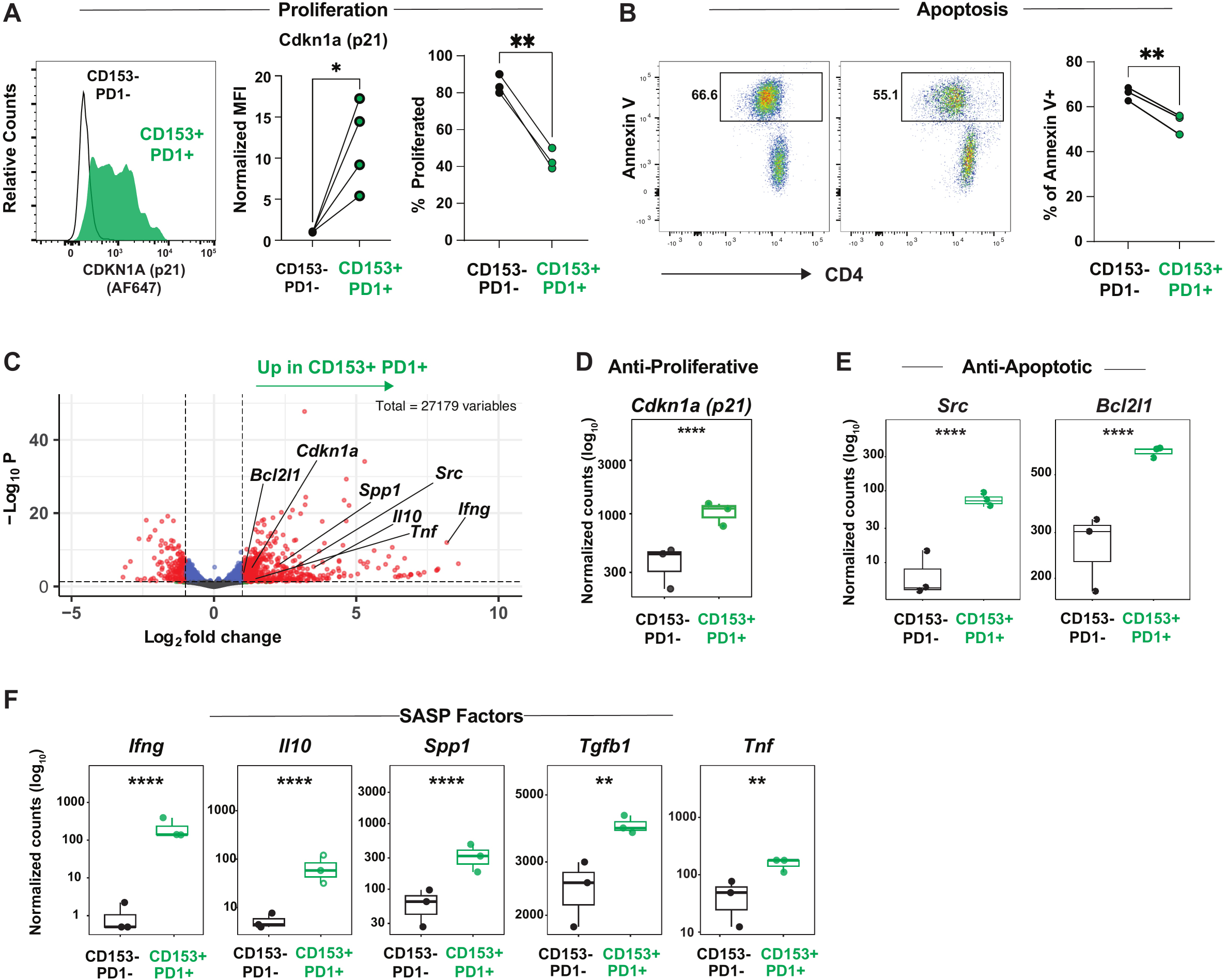
CD153+ CD4+ T cells exhibit a senescence-associated functional phenotype. **A)** Left and center: Representative histograms and normalized mean fluorescence intensity (MFI) quantification by flow cytometry of *Cdknla* expression in *NOD.Aire^GW/+^* peripheral nerve-infiltrating CD153+ PD1+and CD153-PD1- CD44+ CD4+ T cells (*P < 0.05 by paired, two-tailed t test). Right: Percentage of proliferated cells in CD153+ PD1+ or control CD153-PD1- CD4+ T cells sorted from *NOD.WT* spleens (n=3 mice, **P<0.01, paired t-test). **B)** Representative flow plot and percentage of Annexin V+ cells in CD153+ PD1+ and control CD153- PD1- CD4+ T cells from *NOD.WT* spleens (n=2 mice with in vitro replicates, **P<0.01, paired t-test). Pairs represent cells gated from the same mouse. **C)** Volcano plot of upregulated genes by bulk-RNAseq in sorted CD153+ PD1+ and CD153- PD1- T cells from *NOD.Aire^GW/+^* spleens. Padj<0.05, Log2 fold change > 1. **D-F)** Normalized counts for indicated genes by bulk RNAseq of CD153+ PD1+ CD4+ T cells vs. CD153- PD1- CD4+ T cells from *NOD.Aire^GW/+^* mice. **P<0.01, ****P<0.0001, paired t test.

Notably, senescent cells are defined by expression of pro-inflammatory SASP factors, which not only promote chronic inflammation but also spread senescence to neighboring cells, a process known as secondary senescence^36, 37^. Bulk RNAseq of sorted CD153+ PD1+ vs. CD153- PD1- CD4+ T cells demonstrated upregulation of multiple SASP transcripts (*Il10*, *Ifng*, *Tgfb1*, *Spp1*, and *Tnf*) in CD153+ PD1+ CD4+ T cells (**Figure 3C,F**)^15, 38^. Increased SASP factor gene expression was accompanied by significant enrichment of our novel ‘CD4-Old’ core gene signature (**Figure S3B**, left; NES = 2.9, Padj = 9.5e-22), with 37 genes shared between differentially upregulated genes in CD153+ CD4+ PD1+ T cells and the ‘CD4_Old’ signature genes (**Figure S3C)**. Thus, CD153+ PD1+ CD4+ T cells overexpress anti-proliferative, anti-apoptotic, and SASP-associated factors, as well as a core signature of aging-associated features, demonstrating a senescence-associated functional phenotype.

### CD153+ CD4+ T cells exhibit increased autoimmune pathogenic capacity

CD153+ PD1+ CD4+ T cells exhibit multiple functional features of senescence, yet their pathogenic capacity remains unclear. We and others have previously published that CD4+ T cells from spleen and lymph nodes of neuropathic mice are sufficient to transfer autoimmune neuropathy to immunodeficient *NOD.SCID* recipients^7, 8^. Given this, we utilized this system to assess the pathogenic capacity of CD153+ CD4+ T cells, compared total CD4+ T cells (positive control) or CD153- CD4+ (negative control) T cells **(Figure 4A)**. Notably, recipients of CD153+ CD4+ T cells developed overt neuropathy significantly earlier than recipients of CD153- CD4+ T cells **(Figure 4B**, green vs. black line). Consistent with accelerated disease course, histological examination of sciatic nerves revealed increased immune infiltration in recipients of CD153+ CD4+ T cells, compared to CD153- CD4+ T cells (**Figure 4C,D**).

**Figure 4.**
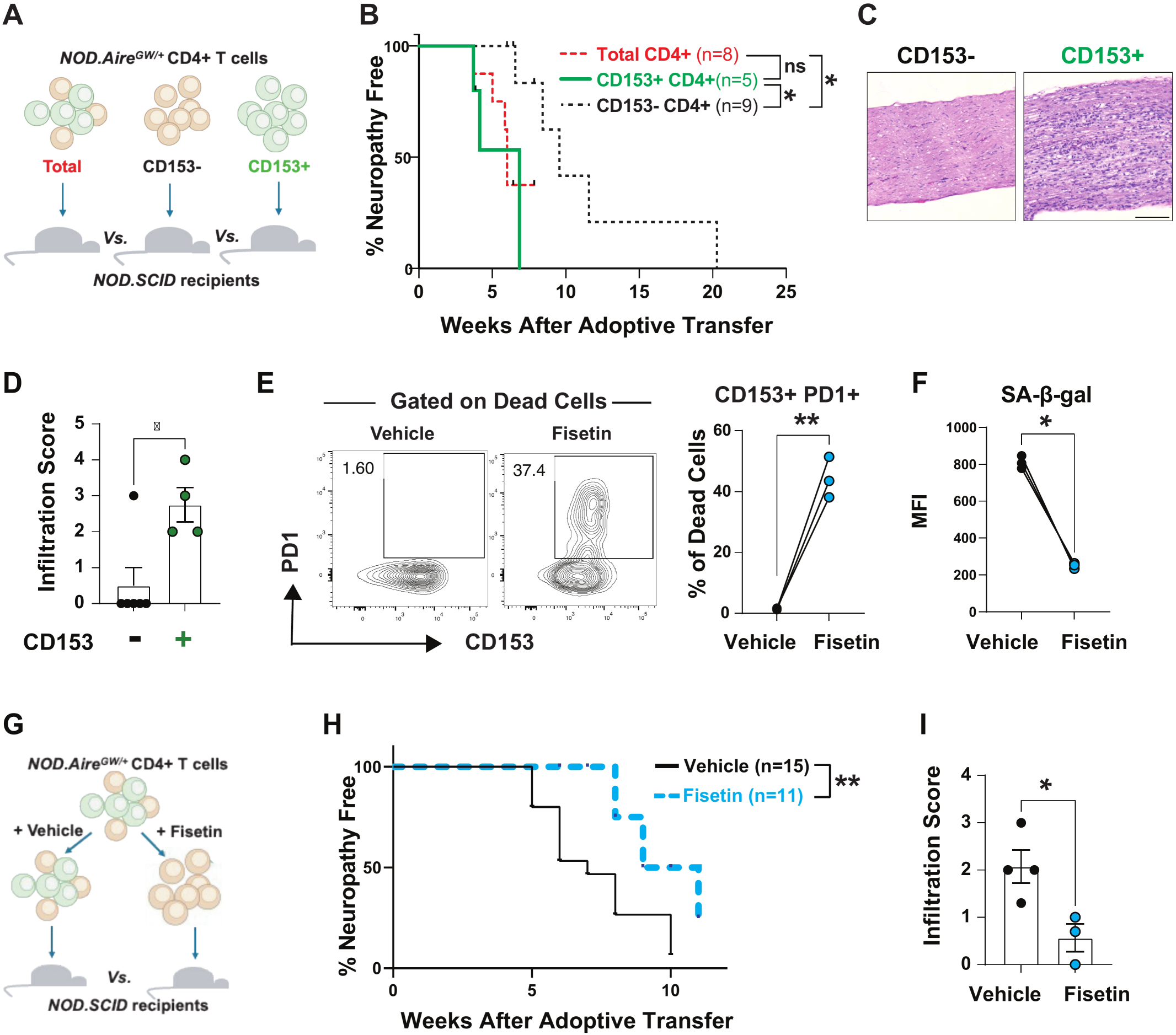
CD153+ CD4+ T cells have enhanced autoimmune capacity. **A)** Scheme for testing the autoimmune capacity of CD153+ CD4+ T cells. **B)** Neuropathy free incidence curve for adoptive tranfer of total, CD153+, and CD153- control CD4+ T cells to recipients (*p-value<0.05, Log-rank test with Bonferroni post-test). **C-D)** H&E staining of sciatic nerves in CD153+ and CD153- control recipients (scale bar=200pm) (C) and scoring of immune cells infiltration (D) (*p-value<0.05, unpaired t-test). **E)** Representative flow cytometry (left) and quantification (right) of CD153+ PD1+ cells among CD4+ T cells expressing dead cell marker following incubation with vehicle or fisetin for 4 days. **F)** SA-p-Gal values of CD4+ T cells from *NOD.Aire^GW/+^* mice treated with fisetin (4ug/ml) or vehicle (DMSO) for 4days in vitro (n=3, p<0.01, paired t-test). **G)** Scheme for adoptive tranfer into *NOD.SCID* recipients of CD4+ T cells treated with fisetin or vehicle. **H)** Neuropathy free incidence curve for adoptive tranfer of CD4+ T cells treated with fisetin or vehicle (** p-value<0.01, Gehan-Breslow-Wilcoxon test). **I)** Immune cell infiltration score in sciatic nerves of *NOD.SCID* recipients of CD4+ T cells treated with fisetin or vehicle (*p-value<0.05, unpaired t-test).

Of note, the presence of suppressive regulatory T cells (Tregs) in the CD153- CD4+ T cell population could account for the delayed neuropathy onset after transfer of this population. However, transfer of CD153+ CD4+ T cells induced neuropathy at the same rate as total CD4+ T cells (positive control), which also contain Tregs **(Figure 4B**, green vs. red line). This finding negates the possibility that a suppressive regulatory T cell population in the CD153- fraction is responsible for the observed delay in neuropathy. Taken together, these findings point to an increased autoimmune capacity within CD153+ CD4+ T cells.

In parallel, we sought to determine the effects of eliminating senescence-associated CD153+ CD4+ T cells on the induction of neuropathy in *NOD.SCID* recipients. To do this, we exploited the senolytic properties of fisetin, a naturally occurring flavonoid that selectively eliminates senescent cells^39, 40^. As expected for a senolytic agent, fisetin treatment resulted in greater death among putative senescent CD153+ PD1+ CD4+T cells **(Figure 4E)** and lowered CD4+ T cell SA-β−Gal activity **(Figure 4F)**. To determine the effects of fisetin on the pathogenic capacity of CD4+ T cells, CD4+ T cells isolated from spleens of neuropathic *NOD.Aire^GW/+^* mice were treated with either vehicle control or fisetin and transferred to immunodeficient hosts (**Figure 4G**). Remarkably, fisetin-treated CD4+ T cells induced neuropathy with delayed kinetics, compared to vehicle-treated controls **(Figure 4H)**, suggesting that elimination of senescent T cells lowers the autoimmune capacity of CD4+ T cells. Indeed, histological examination of sciatic nerves showed reduced immune infiltration in recipients of fisetin-treated CD4+ T cells, compared to vehicle-treated (**Figure 4I**). In support of lower immune infiltration, a significantly decreased number of CD4+ T cells were noted by flow cytometric analysis of sciatic nerves in recipients of fisetin-treated cells (**Figure S3D**). Further, electrophysiologic studies demonstrated significantly decreased Compound Muscle Action Potential (CMAP) latency, and a trend toward increased amplitude and decreased duration, in recipients of fisetin-treated cells (**Figure S3E**). Taken together, these data identify CD153+ CD4+ T cells as a potent autoimmune subset capable of mediating autoimmune disease and highlight the role of the senolytic fisetin to protect from autoimmune nerve injury.

### SASP-associated factors induce senescence in peripheral nerve Schwann cells

Our data thus far demonstrate that senescence associated CD153+ CD4+ T cells exhibit an enhanced capacity for inducing autoimmune peripheral neuropathy (**Figure 4B-D**). It remains unclear, however, the mechanism by which senescence-associated features may lead to increased pathogenicity. CD153+ CD4+ T cells express a number of SASP factors, which have been reported to be capable of inducing secondary senescence in neighboring cells^36, 37^. In infiltrated peripheral nerves, CD4+ T cells are in close proximity to Schwann cells, the primary glial cell type responsible for myelination and repair^8^. Thus, we hypothesized that CD4+ T cell-derived SASP factors induce senescence in neighboring Schwann cells to promote demyelination and neuropathy development (**Figure 5A**).

**Figure 5:**
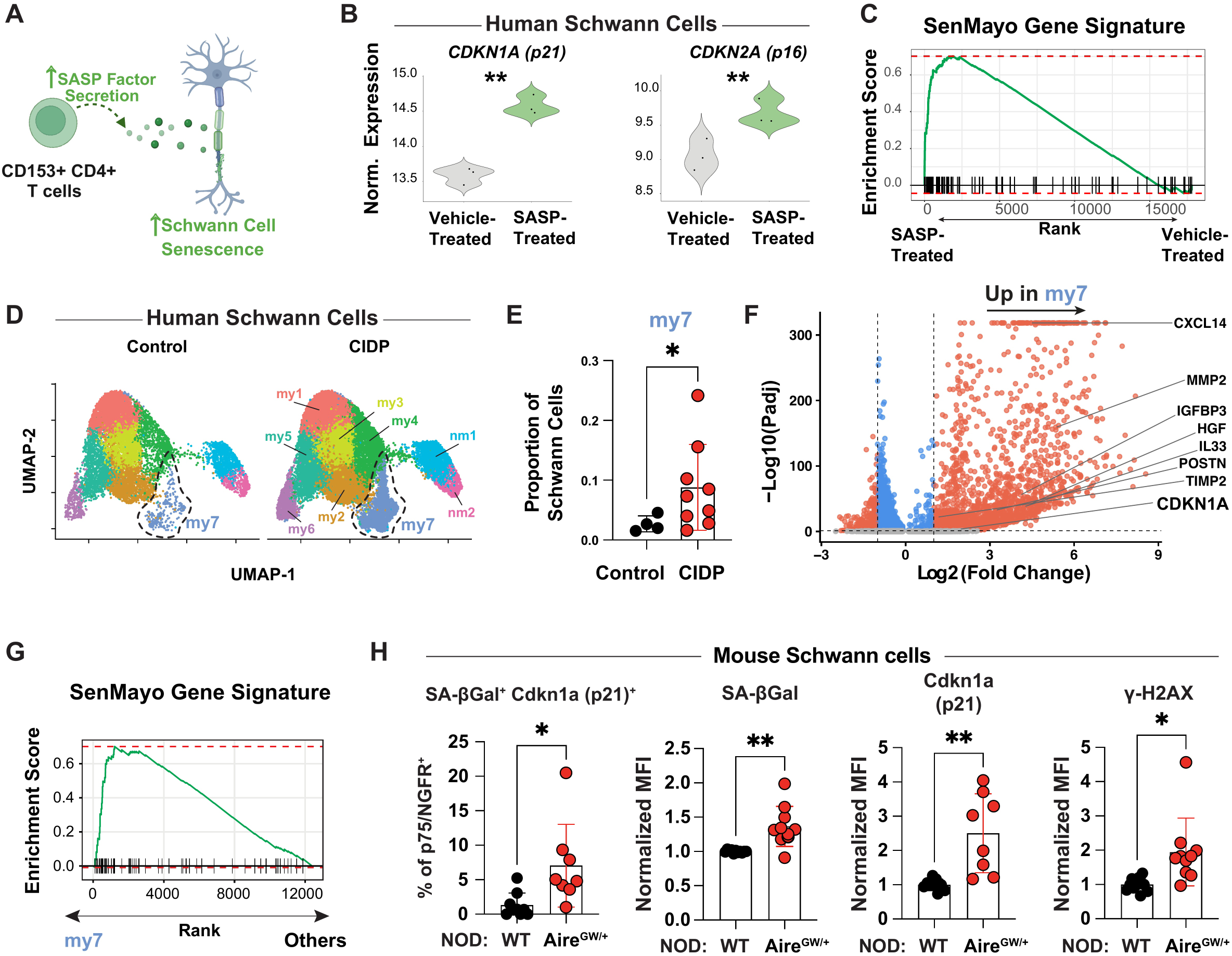
Senescent Schwann cells accumulate in CIDP patients and neuropathic *NOD.AIre^GW/+^* mice. **A)** Model of SASP-mediated Schwann cell senescence induction in autoimmune peripheral neuropathy. **B)** Normalized expression levels of CDKN1A, CDKN2A in stem cell-derived human Schwann with and without SASP (IFNg and TNFa) treatment in vitro. **P<0.01. **C)** GSEAof the senescence-associated SenMayo gene signature in genes upregulated with SASP (IFNg and TNFa) treatment (Padj=1.5e-08). **D)** Uniform manifold approximation and projection of Schwann cells within snRNA-seq data from sural nerve biopsies of CIDP patients (n=9) and age-matched controls (healthy and non-inflammatory neuropathy, n=4). “my” = myelinating, “nm” = non-myelinating. **E)** Comparison of the proportion of Schwann cells belonging to cluster my7 within the snRNA-seq dataset, by patient group (* P<0.05 by two-tailed unpaired t test with Welch’s correction). **F)** Volcano plot of differentially expressed genes upregulated by cluster my7 of the integrated Schwann cell snRNA-seq dataset, relative to all other clusters. Known senescence-associated genes are annotated. **G)** GSEAof the senescence-associated SenMayo gene signature upregulated by cluster my7 relative to all other clusters (****Padj < 0.0001 by fGSEA). **H)** Quantification of senescent Schwann cells expressing indicated markers within the sciatic nerve of *NOD.Aire^GW/+^* and age-matched NOD.WT mice (* P < 0.05, **P<0.01 by two-tailed unpaired t test with Welch’s correction).

To test this possibility, we set up an *in vitro* cell culture system in which CD4+ T cell-derived SASP factors were cultured with human Schwann cells. We focused on two SASP factors, IFNγ and TNFα, that are differentially over-expressed by senescence-associated CD153+ PD-1+ CD4+ T cells (**Figure 2B, 3E**). In this culture system, we also utilized stem cell-derived human Schwann cells, which faithfully recapitulate expression of multiple lineage-specific features including myelination of peripheral nerve axons^41^. Proteomic analysis of these cells confirmed expression of canonical Schwann-cell markers SOX10, MPZ, and MBP (**Figure S4A, Supplementary Table 5**). Bulk RNA sequencing of Schwann cells after incubation with SASP factors (IFNγ/TNFα) vs. vehicle control revealed upregulation of key senescence-associated features, including *CDKN1A* (p21), *CDKN2A* (p16), and *IL6*^22, 42, 43^ (**Figure 5B, S4B; Supplementary Table 6,** Padj<0.05, Log2FC>0.5). Additionally, treated Schwann cells differentially upregulated transcripts that were enriched for the SenMayo senescence gene signature (**Figure 5C**), demonstrating an increase in expression of genes associated with generalized cellular senescence. Thus, SASP factors associated with CD153+ CD4+ T cells are capable of upregulating senescence features in stem cell-derived human Schwann cells.

Motivated by these findings, we next sought to determine whether Schwann cells in peripheral nerves of CIDP patients exhibit features of senescence. To address this, we analyzed a published single-nuclei RNA-sequencing dataset generated from sural nerve samples from CIDP patients (n=9) and age-matched controls (n=2 patients with non-inflammatory neuropathy, n=2 healthy controls)^44^. Sub-clustering analysis of Schwann cells identified nine subsets of myelinating (my1-my7) and non-myelinating (nm1-nm2) Schwann cells (**Figure 5D, Supplementary Table 7**). Of the nine Schwann cell subclusters, cluster my7 was the only population that was uniquely expanded in patients with CIDP (**Figure 5E, S4D-L).**

Remarkably, differential gene expression analysis revealed that relative to other clusters, cluster my7 upregulated the senescence-associated marker *CDKN1A* (p21)^22, 42, 43^ (**Figure 5F, Figure S4C**). Additionally, several SASP factors were upregulated by cluster my7, including *CXCL14*, *IGFBP3*, *POSTN*, *MMP2*, *TIMP2*, *IL33*, and *HGF* (**Figure 5F**). Of particular interest are *CXCL14* and *IGFBP3*, since these SASP members are induced by *CDKN1A* (p21), participate in the recruitment of immune cells^45^, and can induce senescence in multiple cell types^46, 47^. Furthermore, *POSTN* (periostin) has been identified as a pathogenic Schwann cell derived signal which promotes macrophage recruitment in autoimmune peripheral neuropathy^48,49^. Finally, GSEA revealed significant enrichment of SenMayo transcripts within cluster my7 relative to all others (**Figure 5G**). Taken together, these findings suggest that Schwann cells with a senescent phenotype are over-represented in peripheral nerves of CIDP patients.

In parallel, we used flow cytometry of Schwann cells (viable, CD45- p75/NGFR+) from neuropathic *NOD.Aire^GW/+^* and age-matched *NOD.WT* mice to compare senescence marker expression. *NOD.Aire^GW/+^*mice possessed a higher proportion of Schwann cells expressing the senescence marker *Cdkn1a* (p21) and demonstrating SA-β-gal activity (**Figure 5H, left**), relative to *NOD.WT* mice. Furthermore, increased SA-β-gal activity and higher *Cdkn1a* (p21) expression was seen in *NOD.Aire^GW/+^* Schwann cells (**Figure 5H, center panels**). γ-H2AX, a marker of DNA damage associated with cellular senescence was also upregulated by *NOD.Aire^GW/+^* Schwann cells, relative to *NOD.WT* controls (**Figure 5H, right**)^50^. Together, our data point to acquisition of senescence features by Schwann cells in both CIDP patients and mice with CIDP-like disease.

### Senotherapeutic modification of the T cell SASP protects against neuropathy development

Given the urgent clinical need for more effective therapies in patients with CIDP and GBS, we sought to evaluate senotherapies for potential use in CIDP and GBS patients. Our data highlight the role of immunosenescent CD4+ T cells in autoimmune neuropathy and the ability to attenuate autoimmune nerve injury with immunosenescent CD4+ T cell elimination using senolytic agents like fisetin (**Figure 4H,I)**. Because the safety profile for fisetin in humans is relatively limited^51^, however, we turned to FDA approved medications with senotherapeutic properties.

JAK inhibitors are FDA approved medications with senomorphic activities, altering senescent cells through dampening production of SASP-associated factors^52, 53^. While cytokine signaling through JAK/STAT pathways are associated with inflammation, recent studies have also linked JAK/STAT signaling with cellular senescence^53, 54, 55, 56^, and JAK inhibition has shown efficacy in modulating senescence and aging-associated morbidities^54^. Given this, as well as the success of JAK inhibitors in the treatment of other autoimmune diseases^53, 54, 55, 56^, we focused on the potential of the JAK inhibitor, ruxolitinib, as a therapy for CIDP.

Notably, JAK/STAT signaling pathways (e.g., IL-2 signaling, IL-21 signaling, IL-15 signaling, JAK-STAT signaling after IL-12 stimulation) were increased in CD4+ T cells of CIDP patients compared to controls (**Figure S5A**). Additionally, JAK/STAT signaling pathways (e.g., IL-35 signaling, IL-2 signaling, JAK-STAT signaling pathway) were also more prominent in CD153+ PD1+ compared to CD153- PD1- CD4+ T cells from *NOD.Aire^GW/+^* mice (**Figure S5B**). Upregulation of these JAK-STAT mediated pathways in CIDP patient samples and a CIDP mouse model provided additional rationale for considering the efficacy of ruxolitinib in CIDP.

To determine the effects of ruxolitinib on CD4+ T cell senescence, we first treated sorted senescence-associated CD153+ PD1+ CD4+ T cells from *NOD.Aire^GW/+^* mice with ruxolitinib *in vitro.* Compared to vehicle control-treated cells, ruxolitinib treatment reduced expression of SASP-associated cytokines IFN-γ, IL-10, and IL-21 (**Figure 6B**), demonstrating the SASP-suppressing effects of ruxolitinib treatment. Given this, we tested the effects of oral ruxolitinib vs. placebo in *NOD.Aire^GW/+^* mice beginning at 12 weeks of age on neuropathy development. While 70% of mice receiving placebo (n=10) developed neuropathy by 30 weeks of age, mice receiving ruxolitinib therapy were completely protected from clinical signs of neuropathy during this same time frame (**Figure 6C**). Additionally, oral ruxolitinib treatment completely protected against neuropathy in an adoptive transfer model of autoimmune peripheral neuropathy (**Figure S5C)**. Thus, ruxolitinib provides robust *in vivo* protection from neuropathy development, in both a spontaneous model as well as an adoptive transfer model of PNS autoimmunity.

**Figure 6:**
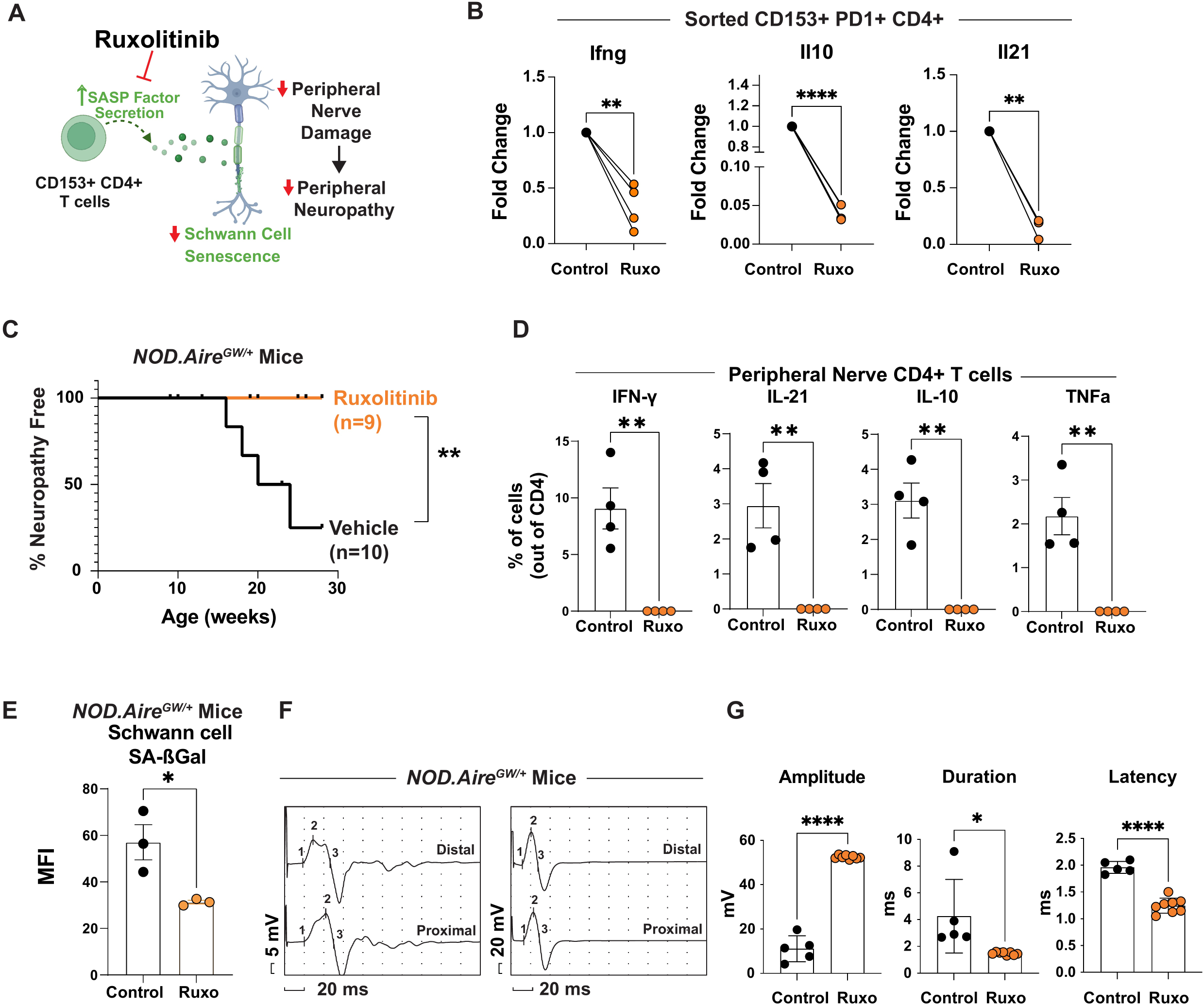
Senomorphic agent Ruxolitinib protects against neuropathy by decreasing CD4+ T cell SASP factor production. **A)** Model of ruxolitinib mechanism of action in protecting against neuropathy. **B)** Expression of SASP transcripts Ifng, 1110, and 1121 by sorted, neuropathic NOD.Aire^GW/+^ CD4^+^CD153^+^ cells treated with ruxolitinib (Ruxo) or vehicle (Control). RT-qPCR data normalized to vehicle-treated CD153^+^ cells (**P < 0.01, ****P < 0.0001 by paired, two-tailed t test). **C)** Kaplan-Meyer survival curve of neuropathy incidence in NOD.Aire^GW/+^ mice treated with ruxolitinib or vehicle control (** P < 0.01, by LogRank test). **D)** Flow cytometry quantification of cytokine-producing CD4+ cells in peripheral nerves of NOD.Aire^GW/+^ mice treated with ruxolitinib or vehicle (** P < 0.01 by two-tailed unpaired t test). **E)** Mean Fluorescent Intensity (MFI) of SA-bGal activity in Schwann cells of NOD.Aire^GW/+^ mice treated with ruxolitinib and vehicle for 1 month. **F)** Representative compound muscle action potentials (CMAP) of NOD.Aire^GW/+^ mice treated with ruxolitinib or vehicle control. **G)** Quantified CMAP parameters comparing ruxolitinib and vehicle treated NOD.Aire^GW/+^ mice (* P < 0.05, **** P<0.0001 by two-tailed unpaired t test).

Within the peripheral nerves of *NOD.Aire^GW/+^* mice, ruxolitinib treatment was associated with decreased CD4+ T cell production of multiple SASP factors (IFN-γ, IL-21, IL-10, and TNFa) (**Figure 6D**), compared to vehicle control treatment, providing support for ruxolitinib’s function in decreasing senescent T cell SASP. Given our model that T cell production of SASP factors induces secondary senescence in nearby Schwann cells (**Figure 6A**), we next sought to determine the effect of ruxolitinib on Schwann cell senescence. Indeed, ruxolitinib treatment was associated with lower Schwann cell SA-β−Gal activity (**Figure 6E**), highlighting the impact of ruxolitinib in preventing senescence induction in Schwann cells. To assess the effects of *in vivo* ruxolitinib treatment on peripheral nerve damage, we performed motor nerve conduction studies on *NOD.Aire^GW/+^* mice that received either ruxolitinib or vehicle control. These studies revealed significant improvement in compound muscle action potential (CMAP), with significantly increased CMAP amplitude and significantly decreased CMAP duration and latency in ruxolitinib-treated mice (**Figure 6F, G**). These findings provide further evidence that ruxolitinib protects *NOD.Aire^GW/+^* mice from peripheral nerve damage and demyelination at the tissue level. In summary, our findings identify a clinically-approved JAK inhibitor with senomorphic action as a potential near-term strategy to target highly pathogenic senescence-associated T cells and protect against autoimmune peripheral neuropathies.

## Discussion

Recent advances have extended the human lifespan and contributed to an unprecedented increase in the older population^57^. However, much work remains to improve healthspan. Aging compromises multiple aspects of immune function, including mechanisms of self-tolerance^1^. How age-associated changes predisposes to certain autoimmune diseases, however, remains unclear. We show here that circulating CD4+T cells from patients with CIDP, a condition associated with increased age, exhibit multiple immunosenescence features. In the *NOD.Aire^GW/+^* mouse model of CIDP, a population of senescence-associated CD153+ CD4+ T cells accumulate in peripheral nerves and demonstrate an increased capacity for inducing autoimmune peripheral neuropathy. Together, these findings demonstrate a critical role for senescence-associated CD4+ T cells in the pathogenesis of PNS autoimmunity.

Studying age-associated changes in T cells is complicated by the lack of uniform biomarkers to identify senescent T cells. While *Cd153* has been identified as a senescence marker in some T cell populations^15, 16^, other senescence-associated markers [e.g., *Klrg1*^58, 59^, *Cdkn2a (p16)*^60^, *Tnfa*^38^] have also been reported. Moreover, there is overlap in cytokines produced as part of the SASP by senescent T cells and those produced by T cells with activation. To address this, we exploited the Tabula Muris Senis, a single-cell atlas of the aging mouse^61^, to generate a ‘CD4-Old’ core gene signature. Because the Tabula Muris Senis dataset was built using an autoimmune resistant strain of mice (C57BL/6), we reasoned that biomarkers associated with CD4+ T cells in old mice in this dataset would not be confounded by markers upregulated with autoimmune responses. This ‘CD4-Old’ signature supported our identification of a CD4+ T cell subset that exhibits features of age-associated senescence in our mouse model of CIDP and may be a resource for future studies seeking to identify senescent CD4+ T cells beyond CIDP.

Functional analyses of CD153+ CD4+ T cells demonstrated multiple features of senescence, including decreased proliferation, resistance to apoptosis, and expression of a pro-inflammatory SASP. A number of these SASP factors (*Il10*, *Ifng*, *Tgfb1, Tnf*) have been linked to the pathogenesis of autoimmune peripheral neuropathy in mice and humans^48, 62, 63, 64, 65^. Importantly, SASP factors foster a pathologic microenvironment not only by promoting inflammation, but also by spreading senescence to neighboring cells^36, 37^. Indeed, we found that senescence features are exhibited by peripheral nerve Schwann cells in CIDP patient samples and neuropathic *NOD.Aire^GW/+^*mice, and senescence could be induced in Schwann cells after incubation with SASP factors produced by senescent T cells. Taken together, these data suggest a model in which SASP factors produced by CD4+ T cells spread senescence to neighboring Schwann cells.

While CD153 expression is a common feature of senescence-associated CD4+ T cells in CIDP patients and *NOD.Aire^GW/+^* mice, whether CD153 is merely a marker of senescent T cells or plays a role in the induction of senescence is unclear. Prior work has reported that CD153 engagement with TCR induces pathogenic senescence in T cells^34^. Moreover, CD153 can also interact with CD30 on B lymphocytes, mediating development of germinal center B lymphocytes and augmenting autoimmunity in a mouse model of lupus^34^. Of note, B lymphocytes are dispensable for *NOD.Aire^GW/+^* neuropathy^62^, suggesting that any CD153 mediated effects would be independent of its effects of B cells in this model. Future studies to knockout CD153 or block its interactions with CD30 may be useful in delineating CD153’s role in autoreactive T cells.

Understanding pathological senescence presents multiple therapeutic opportunities for age-associated autoimmunity. Senotherapies that deplete or modulate senescent cells have shown efficacy in a number of age-related diseases^66^ but have not yet been tested in autoimmune peripheral neuropathy. Our data indicate that depletion of senescent CD4+ T cells with a senolytic agent (fisetin) reduces the capacity of CD4+ T cells to induce neuropathy upon transfer into an immunodeficient host. Additionally, we demonstrate that treatment of *NOD.Aire^GW/+^* mice with ruxolitinib, an FDA-approved, orally available senomorphic therapy, prevents neuropathy development. Thus, our findings provide preclinical evidence for the use of senotherapies as a novel preventative approach for age-associated autoimmune conditions.

## Materials and Methods

### Human sample collection

Approval to identify and consent patients to collect peripheral blood samples for this study was granted by the UCLA Institutional Review Board (IRB-19-0032: Mechanisms of Autoimmunity). Patients with CIDP were identified by a board-certified neurologist and consented to provide peripheral blood samples before initiation of immunomodulatory therapies. Peripheral blood mononuclear cells (PBMCs) were isolated using Ficoll-Paque PLUS (Cytiva #17144002) density gradient centrifugation, then stored in liquid nitrogen in freezing media (fetal bovine serum with 10% DMSO). Samples from patients who later responded to immunomodulatory therapy consistent with CIDP were selected for flow cytometry and single cell RNA sequencing studies. Age-matched healthy control PBMC samples were purchased from BioChemed Services. The demographic information of CIDP patients and healthy controls is provided in **Supplementary Table 9**.

### Mice

*NOD.Aire^GW/+^*, *NOD.WT*, and *NOD.Prkdc^SCID^*mice were bred and housed in a pathogen-free barrier facility maintained by the UCLA Division of Laboratory Animal Medicine. Breeding and experimental protocols were approved by the UCLA Animal Research Committee and Institutional Animal Care and Use Committee (protocols ARC-2018-082 and ARC-2018-087).

### *In vivo* ruxolitinib administration

Mice were provided *ad libitum* access to medicated (1 g ruxolitinib/kg food gel) or vehicle-control food gels starting at 9 weeks of age. Ruxolitinib (MedChem Express, #HY-50856) was added to Nutra-Gel Diet dry mix (Bio-Serv, #F5769-KIT, Flemington, NJ), and food gels were replaced daily for twenty weeks.

### Flow cytometry

After thawing and resting human PBMC samples overnight, samples were cultured in complete RPMI and stimulated with anti-human CD3/CD28 Dynabeads (1:1 cell to bead ratio, Thermo Fisher Scientific #11131D) for two days. Dynabeads were then removed, and cells were resuspended in fresh complete media with human IL-2 (20ng/mL) and cultured for an additional 13 days. Complete RPMI with IL-2 was replaced on day 7 post-stimulation. Following 13-day culture with IL-2, samples were stained with SA-βGal, viability dye, and surface markers (**Supplementary Table 8**).

For flow cytometric analysis of mouse Schwann cells and nerve-infiltrating senescence-associated T cells, single cell suspensions were prepared by enzymatic and mechanical digestion. Briefly, sciatic or brachial nerves were cut into fine pieces while in digestion buffer (1mg/mL collagenase IV in PBS with 1% FBS), then incubated at 37C for 20 minutes, vortexing every 5 minutes. Following trituration with a 20 gauge needle, samples were incubated for an additional 10 minutes at 37C, triturated, then filtered with a 40 micron cell strainer. Cells were stained with surface and intracellular antibodies as indicated in (**Supplementary Table 8**).

For SA-bGal staining, human or mice cells were incubated with bafilomycin A1 (Dojindo Molecular Technologies #SG04) in complete RPMI for 1 hour at 37C with 5% CO2. Samples were then stained with SPiDER-βGal (Dojindo Molecular Technologies #SG04) in complete media with bafilomycin A1 for an additional 30 minutes at 37C with 5% CO2. After SPiDER-βGal staining, samples were stained with Fixable Viability Dye and surface and intracellular markers.

Cells were fixed and permeabilized for intracellular staining using the eBioscience FOXP3/Transcription Factor Staining Buffer Set (Thermo Fisher Scientific #00-5523-00) or Fixation (Invitrogen, GAS001S) and Permeabilization medium (Invitrogen, GAS002S). Sample fluorescence was measured using an Invitrogen Attune NxT flow cytometer (Thermo Fisher Scientific), or BD LSRFortessa, then analyzed using FlowJo (version 10.10.0, Becton Dickinson and Co.).

### Cell sorting

For bulk-RNA seq experiment, adoptive transfer, and in vitro treatment of senescence-associated T cells, splenic CD4+ cells were isolated by magnetic separation (STEMCELL Technologies, Mouse EasySep CD4 T cell isolation kit #19852), followed by fluorescent isolation of CD153+PD1+ and CD153-PD1- CD4+CD44+ cells by a BD FACS Aria III (**Supplementary Table 8**). For adoptive transfer and *in vitro* treatment with ruxolitinib experiments, CD153-expressing cells were stimulated and isolated using the protocol above, but were not stained with anti-CD4, anti-CD44, or anti-PD1 to avoid changes in T cell phenotype due to agonism/blockade.

### Proliferation and Apoptosis of Senescent associated T cells

CD153+PD1+ and CD153-PD1- splenocytes from NOD.WT mice (17-26 weeks), were sorted, labeled with CellTrace™ Violet Cell Proliferation Kit (Invitrogen, C34557), and cultured for 3 days in complete T cell medium (DMEM, 10% FBS, PenStrep, B-Mercaptoethanol). Following culture, cells were stained with surface markers and Annexin V (**Supplementary Table 8**) before acquisition on a BD Fortessa flow cytometer.

### Adoptive transfers, histology, and Electromyography (EMG)

Sorted CD153+ CD4+ and CD153- CD4+ T cells were resuspended in sterile PBS (1×10^5^ cells/100uL). 6-10 week old *NOD.Prkdc^SCID^* recipients were anesthetized with isoflurane prior to retro-orbital injection of 1×10^5^ cells. Four weeks after adoptive transfer, recipients were checked daily for clinical signs of neuropathy, as previously described^48^. Sciatic nerve sections were collected from paired samples (CD153+ and CD153- recipients) upon the development of neuropathy, fixed with 10% neutral-buffered formalin, then paraffin embedded and stained with hematoxylin and eosin by the UCLA Translational Pathology Core Laboratory (TPCL). Nerve infiltration (0: no infiltration, 1: early infiltration, 2: mild infiltration, 3: moderate infiltration, 4: severe infiltration) was scored by three blinded observers, then averaged.

To assess the effect of fisetin in neuropathy, CD4+ cells isolated from spleen of neuropathic mice and treated with 4μg/ml Fisetin (MedChem, HY-N0182) or DMSO as vehicle for 4days. Cells were collected and equal number of cells injected to_*NOD.Prkdc^SCID^* recipients retro-orbitally. To assess the effect of Ruxolitinib treatment in an adoptive transfer model, splenocytes were isolated from a neuropathic *NOD.Aire^GW/+^* mice and treated for 4 days in vitro (1ug/ml aCD3 and 1ug/ml aCD28). Then cells transferred to *NOD.Prkdc^SCID^* recipient (r.o) and mice were randomly assigned to Ruxolitinib and Vehicle cohorts.

EMG was performed using a TECA Synergy N2 EMG machine as previously described^32^. Compound muscle action potentials (CMAPs) were recorded following the stimulation of the sciatic nerve for 0.1 ms duration at 1 Hz frequency and 20 mA intensity stimulus, with the low-pass filter set to 20 Hz and the high-pass filter to 10 kHz.

### Single-cell RNA sequencing of peripheral human T cells

PBMC samples were thawed and rested overnight in complete RPMI. T cells were then isolated from rested PBMCs using magnetic negative selection (Miltenyi Biotec #130-096-535) and sent to the UCLA Technology Center for Genomics and Bioinformatics core for Next-GEM 5’GEX + TCR v2 library construction (10X Genomics) and Illumina NovaSeq X Plus sequencing. 10X Genomics CellRanger Count (v8.0.0) was used to align reads (human genome (GRCh38) 2024-A) and aggregate transcript counts per cell, followed by Seurat analysis (version 5.3.0). Briefly, cells with < 200 features, > 2500 features, or > 5% mitochondrial DNA were removed. Highly variable genes were identified with the mvp selection method of the Seurat FindVariableFeatures function. TCR (alpha/beta chains) and BCR (heavy/light chains)-encoding genes were excluded from the set of highly variable features. Data were scaled using default parameters, with regression for total RNA per cell (nCount_RNA). Principal component analysis (PCA) was performed using default parameters. 20 dimensions and a resolution of 0.5 were used for nearest-neighbor calculation, clustering and uniform manifold approximation and projection (UMAP). After integration with healthy control samples, CD4 T cells were subset by excluding CD8-expressing cells (CD8A expression < 0.25) and selecting cells expressing CD4 (CD4 expression > 0.25). In total, 2026 CD4+ T cells in CIDP and 915 CD45+ T cells from controls analyzed further. The number of cells per individual is listed in **Supplementary Table 10.** 40 dimensions and a resolution of 0.5 were used for subsequent reclustering. R code to reproduce data visualization, differential gene expression, and cluster proportion analyses is available on Code Ocean.

### Mouse single cell RNA-sequencing analysis

For analysis of sciatic nerve infiltrating senescence-associated T cells within the *NOD.WT* and *NOD.Aire^GW/+^* models, previously published nerve-infiltrating CD45+ scRNA-seq datasets were analyzed. Clusters expressing Cd3e (T cells) were subset from an existing integrated dataset and preprocessed using the default parameters for Seurat (version 5.3.0) normalization, variable feature identification, scaling, and principal component analysis. 30 dimensions and a resolution of 0.5 were used for clustering and UMAP of the T cell subclusters. CD4 T cells were then subset from the T cell dataset through exclusion of clusters expressing *Cd8a* and *Tcrg-V5*. In total, 5204 CD4+ T cells from WT and 23316 CD4+ T cells from *NOD.Aire^GW/+^* mice analyzed. The subset of CD4 T cells was then subject to normalization, variable feature identification, scaling, and principal component analysis using the default Seurat parameters. 25 dimensions and a resolution of 0.4 were used for clustering and UMAP. R code to reproduce data visualization, differential gene expression, and cluster proportion analyses is available on Code Ocean. The number of cells per mouse is listed in **Supplementary Table 10.** The ’Cd4Old’ core gene signature was generated from the Tabula Muris Senis dataset^22^. Among differentially expressed genes between CD4+ T cells from old and young mice in Tabula Senis Muris^22^, genes up-regulated in old (padj<0.05, Log2FC>1) were included in the signature.

### Bulk RNA-seq analysis

RNA was isolated from CD153+ CD4+ and CD153- CD4+ T cells using a Zymo Quick-RNA MicroPrep kit (Zymo Research #R1051). Subsequent mRNA purification and cDNA synthesis were performed using the ABclonal Stranded mRNA-seq Lib Prep Kit for Illumina (ABclonal, RK20301), followed by Illumina NovaSeq X Plus sequencing. Cutadapt (version 5.1) was used to trim adapter sequences and poly-G artifacts from NovaSeq sequencing. Trimmed reads were then aligned to the mm10 mouse genome using HISAT2 (version 2.2.1). Gene expression counts were calculated using featureCounts (version 2.1.1). DESeq2 (version 1.44.0) was used to normalize raw counts, perform blinded principal component analysis, and to perform differential gene expression testing. Differentially expressed genes were ranked by Wald significance and used to run gene set enrichment analysis with fGSEA (version 1.30.0). R code to reproduce data visualization, differential gene expression, and gene signature enrichment analyses is available on Code Ocean.

### Human snRNA-seq analysis

Gene expression matrices (Cellbender H5 format) were downloaded from GEO (GSE285983) and imported to R using the hdf5r package (version 1.3.12). After conversion to sparce matrices with the Matrix package (version 1.7-3), the raw counts were used to construct Seurat (version 5.3.0) objects. Cells with < 200 features, > 9000 features, or > 5% mitochondrial DNA were removed. Doublets were removed following identification with scDblFinder (version 1.18.0). To improve memory efficiency, BPCells (version 0.3.0) was used to store counts data on-disk. Default parameters were used for Seurat normalization, variable feature selection, scaling, and PCA. 30 dimensions were used for clustering and UMAP. Schwann cells were subset from each object by expression of *SOX10, S100B, MBP, PRX, CDH1, SLC36A2, L1CAM, CDH2*, and *CHL1*. Schwann cell subsets from CIDP (27333 cells) and control samples (10378 cells) were merged and subject to Seurat normalization, variable feature selection, scaling, PCA, and RPCA integration using default parameters. 20 dimensions were used for clustering and UMAP analyses. R code to reproduce data visualization, differential gene expression, gene signature enrichment, and cluster proportion analyses is available on Code Ocean. The number of cells per individual is listed in **Supplementary Table 10.**

### *In vitro* Schwann cell stimulation

One day prior to stimulation, embryonic stem cell-derived Schwann cells (ESC-SCs) (a gift from Dr. Faranak Fattahi)^41^ were seeded onto a 96 well TC-treated plate that had been coated with Poly-L-Ornithine (Sigma Aldrich Cat# P3655-10MG), fibronectin (Thermo Fischer Cat# 33016-015), and laminin (R&D systems Cat# 3400-010-02). Cells were cultured in Schwann cell media (Sciencell #1701) supplemented with Schwann cell growth supplement. Cells were treated for twenty-four hours with media only or 100 U/mL of human recombinant IFN-g in addition to 10 ng/mL of human recombinant TNFa. Cells were lysed with RNA lysis buffer and subsequently processed for bulk RNA-sequencing.

### Mass spectrometry proteomics of stem-cell-derived Schwann cells

Samples were lysed in 8 M urea (Promega) in LoBind tubes (Eppendorf), kept on ice, and probe-sonicated 5 × 10 s with 10 s intervals at 10% amplitude. Protein concentration was measured by BCA assay, and ∼100 µg protein per sample was reduced with 10 mM TCEP and alkylated with 40 mM 2-chloroacetamide for 30 min each at 23°C with shaking at 1100 rpm. Urea was diluted eightfold with 100 mM Tris-HCl (pH 8), and proteins were digested overnight (16 h) at 33°C and 1000 rpm with trypsin (Promega) and Lys-C (Wako), each at a 1:100 enzyme:substrate ratio. Samples were acidified with TFA to pH ∼2–3 and desalted using 30 mg HLB 1 cc cartridges (Waters), which were activated with 1 mL 80% ACN/0.1% TFA, equilibrated with 3 × 1 mL 0.1% TFA, loaded with sample, washed with 3 × 1 mL 0.1% TFA, and eluted with 0.8 mL 50% ACN/0.25% formic acid. Peptides were dried by vacuum centrifugation, resuspended in 0.1% formic acid, and analyzed on a timsTOF HT mass spectrometer (Bruker Daltonics) coupled to a Vanquish Neo UHPLC. Mobile phases were 0.1% formic acid in MS-grade water (A) and 0.1% formic acid in MS-grade acetonitrile (B). Peptides were trapped on a PepMap Neo Trap column (5 mm, 100 Å, 5 µm) and separated on an Aurora Elite C18 column (15 cm, 100 Å, 1.5 µm; IonOpticks) maintained at 50°C and coupled to a CaptiveSpray source operated at 1700 V. The gradient was 5–35% B over 37 min at 0.3 µL/min, followed by 35–45% B over 4 min, 45–60% B over 1 min, and 60–95% B over 3 min. Data were acquired in dia-PASEF mode using variable isolation windows in the m/z–ion mobility plane optimized for precursor coverage. MS1 scans covered 100–1700 m/z with a dual-TIMS ramp rate of 9.42 Hz and 100 ms accumulation and ramp times (100% duty cycle), an ion mobility range of 0.63–1.52 V·s/cm², 40 Da isolation windows with 1 Da overlap, and 25 mass steps per 1.06 s cycle. Collision energy decreased linearly from 59 eV at 1/K₀=1.6 V·s/cm² to 20 eV at 1/K₀=0.6 V·s/cm², with MS/MS spectra collected from 251.3–1227.3 m/z. Raw files were processed in Spectronaut (Biognosys) using the directDIA+ (Deep) workflow, with carbamidomethylation of cysteine as a fixed modification and protein N-terminal acetylation and methionine oxidation as variable modifications. Reviewed human protein sequences downloaded from UniProt on October 6, 2023 were used for database searching, with false discovery rates for PSMs, peptides, and protein groups controlled at 1%. Protein abundance was quantified from MS2-level peak areas with Spectronaut cross-run normalization disabled. Quantitative analyses were performed in R (v4.4.1), including inter-run clustering, correlations, principal component analysis, peptide and protein counts, and intensity distributions. Peptides mapping to the same protein were summarized using Tukey’s Median Polish, and MSstats was used for cross-run normalization by median equalization without missing-value imputation.

### *In vitro* T cell treatment with ruxolitinib

After isolation of CD153+ and CD153- CD4+ T cells from stimulated *NOD.Aire^GW/+^* splenocytes, 5×10^4^ cells were seeded per well in a 96-well plate coated with anti-CD3 (1ug/mL in PBS, clone 145-2C11, Thermo Fisher Scientific #14-0031-86) and anti-CD28 (1ug/mL in PBS, clone 37.51, Thermo Fisher Scientific #14-0281-86). Cells were treated with ruxolitinib (10uM, MedChemExpress #HY-50856) or vehicle control (DMSO, VWR #0231-500ML) in complete RPMI for 24 hours, then lysed in RNA Lysis Buffer (Zymo Research #R1060-1-50) for RT-qPCR.

### RT-qPCR

RNA from ruxolitinib/vehicle-treated T cells was isolated using the Zymo Research Quick RNA Isolation Kit protocol (#R1050). cDNA was prepared using the Applied Biosystems High Capacity cDNA Reverse Transcription Kit (Thermo Fisher Scientific #4368813). Murine brachial nerve *Cdkn1a* (p21) transcripts were quantified using the Applied Biosystems PowerUp SYBR Green Master Mix (Thermo Fisher Scientific # A25742) with forward primer 5’-TTGCCAGCAGAATAAAAGGTG-3’ and reverse primer 5’-TTTGCTCCTGTGCGGAAC-3’.

Brachial nerve *Cdkn1a* transcript expression was normalized to *Hprt* (Forward: 5’-TTTCCCTGGTTAAGCAGTACAGCCC-3’, reverse 5’-TGGCCTGTATCCAACACTTCGAGA-3’). T cell *Ifng* (Taqman assay ID: Mm01168134_m1), *Il21* (Taqman assay ID: Mm00517640_m1), *Il10* (Taqman assay ID: Mm00439614_m1), and *Actb* (Mm02619580_m1) transcripts were measured using the Applied Biosystems TaqMan Fast Advanced Master Mix (Thermo Fisher Scientific #4444557). Transcript expression was normalized to beta-actin (*Actb*).

## Supporting information

Supplemental Tabel 1-10

## Data availability

Supporting data for all figures has been made available through FigShare. Raw and processed scRNA-seq (human peripheral T cell) and bulk RNA-seq (mouse CD153+ CD4+ and control CD153- CD4+ T cells) data have been made available on the NCBI Gene Expression Omnibus (GSE315020, GSE314953). Bulk-RNAseq data for human Schwann cells have been submitted to GEO and are currently under curation. Code to reproduce scRNA-seq, snRNA-seq, and bulk RNA-seq analyses has been made available on Code Ocean.

## Acknowledgements

The authors would like to acknowledge the assistance of the UCLA JCCC and BSRB Flow Cytometry Core (flow cytometers, FACS), the UCLA Technology Center for Genomics and Bioinformatics (scRNA-seq, bulk RNA-seq), and the UCLA Translational Pathology Core Laboratory (H&E staining). This research was supported by a National Institute of General Medical Sciences of the National Institutes of Health Maximizing Investigators’ Research Award (MIRA) award no. 1R35GM156893-01.

