## Supplemental Figure 1-5 for "Age-associated Autoimmunity Driven by T cell Immunosenescence"

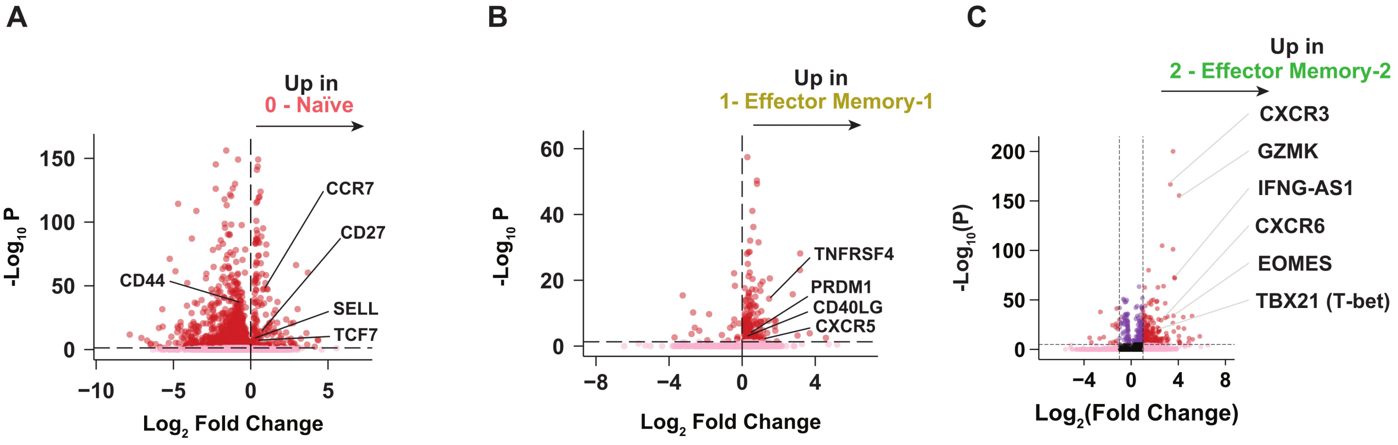


**Supplemental Figure 1: scRNA-seq identifies distinct populations of peripheral CD4 T cells in CIDP and healthy controls. A-C)** Volcano plot of differentially expressed genes upregulated by clusters 0 (A) and 1 (B) and 2 (C) of the integrated healthy control and CIDP peripheral CD4 T cell scRNA-seq dataset, relative to all other clusters.

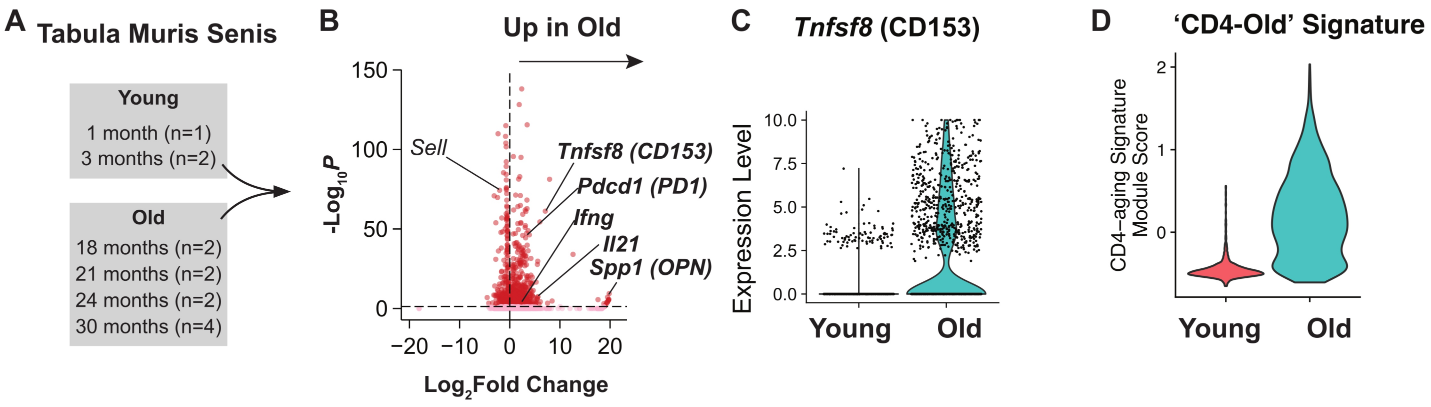


**Figure S2: Generation of a mouse 'CD4Old' gene signature for aging-associated senescent T cells. A)** Schematic of scRNA-seq data analysis using splenic CD4 T cells of young (1-3mo.) and old (18-30mo.) mice from Tabula Muris Senis. **B)** Volcano plot of differentially expressed genes upregulated by old CD4 T cells relative to young CD4 T cells. **C)** Violin plot of Tnfsf8 (CD153) transcript expression within the splenic CD4 T cell scRNA-seq data, split by age group. **D)** Enrichment of 'CD4-Old' signature in CD4+ T cells from old mice in Tabula Muris Senis database.

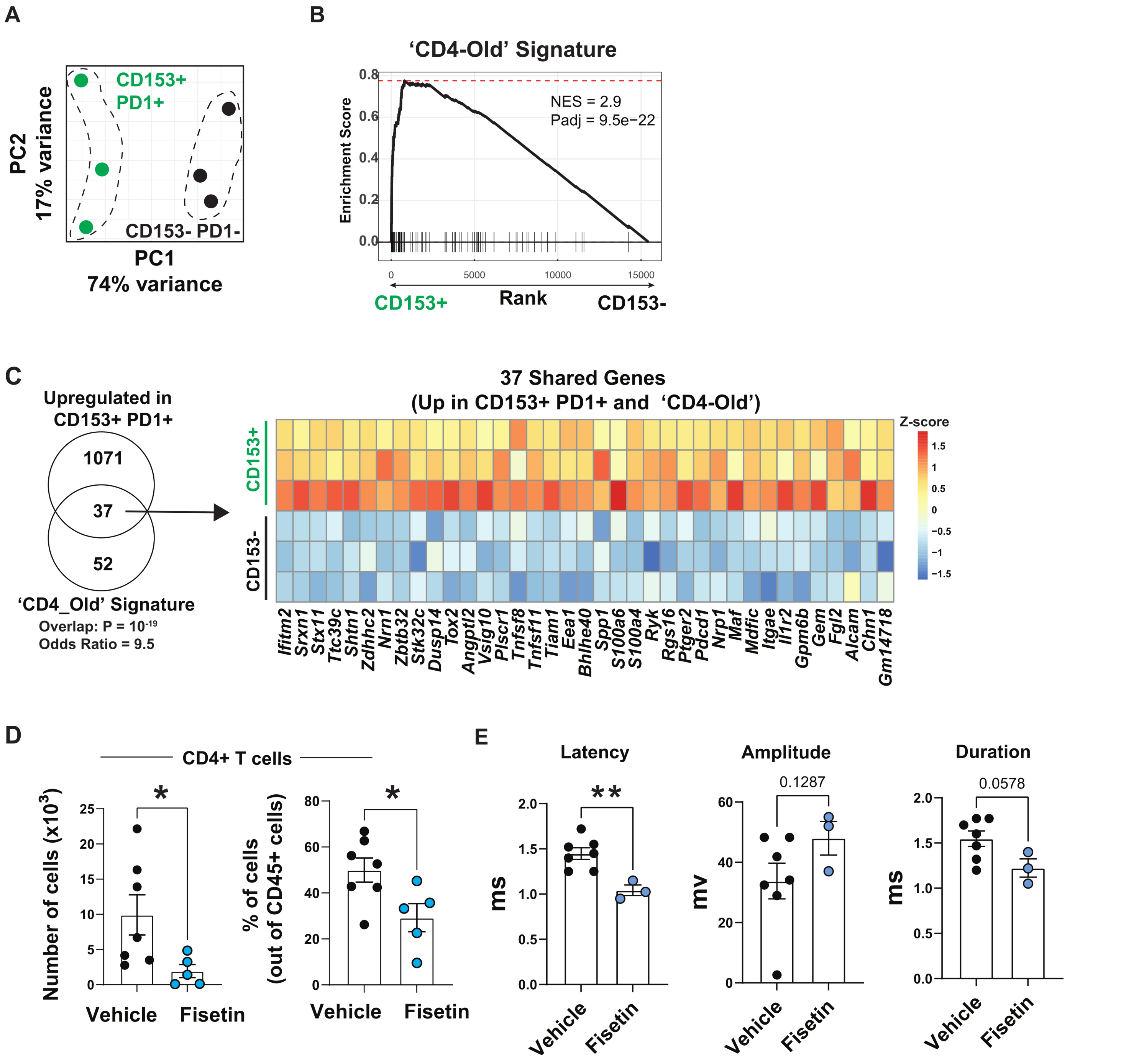


**Figure S3: Characterization of CD153+ CD4+ cells and effects of fisetin on CD4+ T cell pathogenicity. A)** PCA plot of bulk RNAseq of CD153+ PD1+ vs CD153- PD1- CD4+ T cells, sorted from *NOD.Aire*$^{GW/+}$ mice. **B)** GSEA enrichment analysis for "CD4-Old' gene sigature within differentially expressed genes between CD153+ PD1+ vs. CD153- PD1- CD4+ T cells. **C)** Venn diagram and heat map of 37 shared genes that are part of the 'CD4-Old' gene signature and differentially upregulated genes in CD153+ CD4+ T cells. Overlap using Fisher's test: Odds ratio 9.5, P = $10^{-19}$. **D**) Percentage and absolute numbers of infiltrated CD4+ cells in the sciatic nerves of *NOD.SCID* recipients of vehicle vs. fisetin treated CD4+ T cells (n=3-4, *p<0.05, unpaired t-test). **E)** Quantified latency, amplitude, and duration of CMAP of *NOD.SCID* recipients of vehicle vs. fisetin treated CD4+ T cells at 5-7 weeks after transfer (**p<0.01, unpaired t-test).

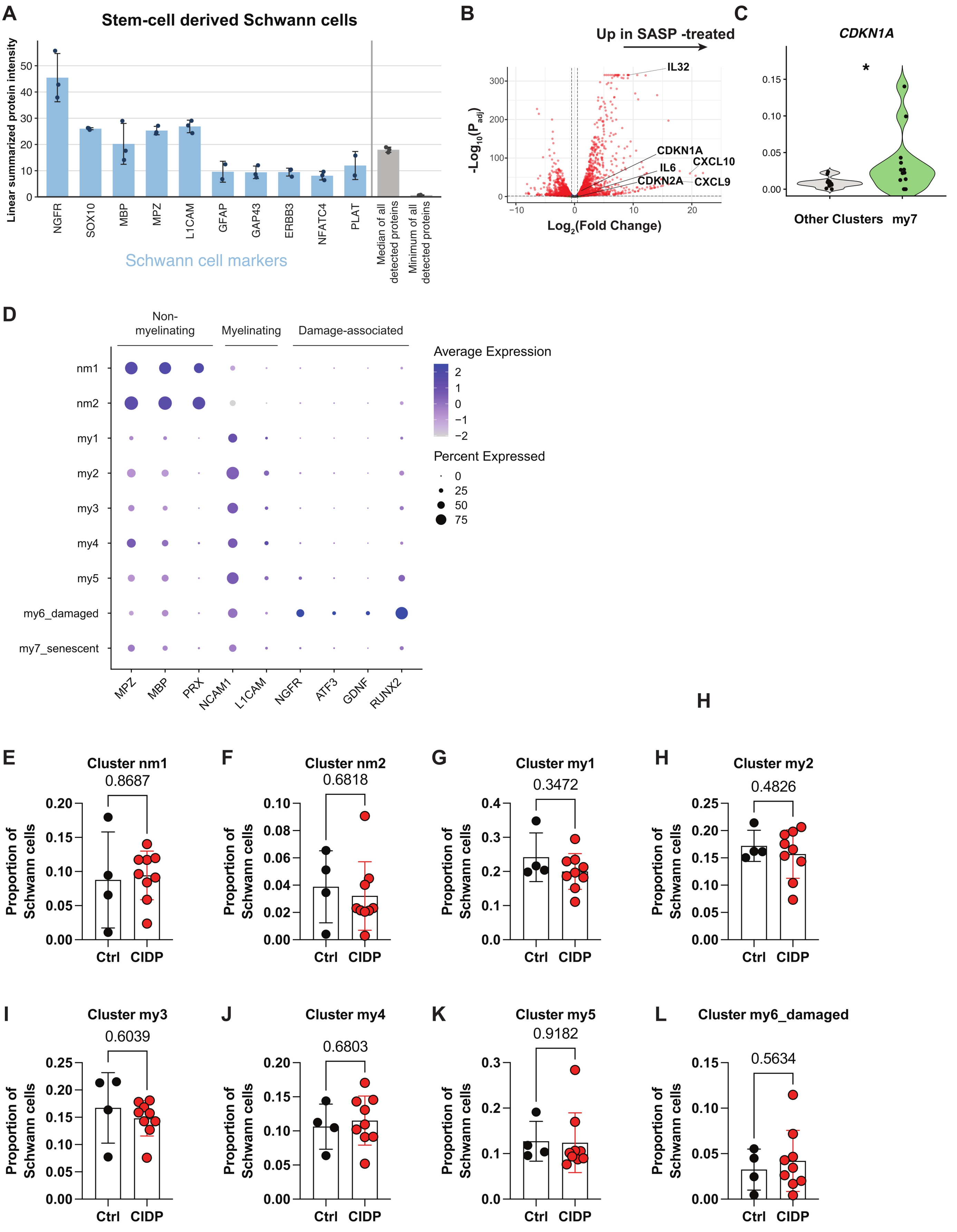


**Figure S4: Stimulation of immortalized human Schwann cells with SASP factors. A)** Proteomic analysis of stem cell-derived Schwann cells. The canonical Schwann-cell markers SOX10 and MPZ were consistently detected across all three iPSC-derived Schwann-cell replicates, ranking in the 67th-70th and 66th-68th percentiles of proteome-wide abundance, respectively, with levels above the median abundance of all detected proteins. **B)** Volcano plot highlighting senescence-associated, upregulated genes following IFNg and TNFa SASP stimulation of stem cell-derived human Schwann cells. **C)** Pseudobulk analysis of CDKN1A (p21) normalized expression in my7 compared to all other clusters.*Padj<0.05 .**D)** Dot plot of Schwann cell transcript expression within the snRNA-seq dataset. **E-L)** Comparison of the proportion of Schwann cells belonging to each cluster within the snRNA-seq dataset, by patient group (by two-tailed unpaired t test with Welch's correction).

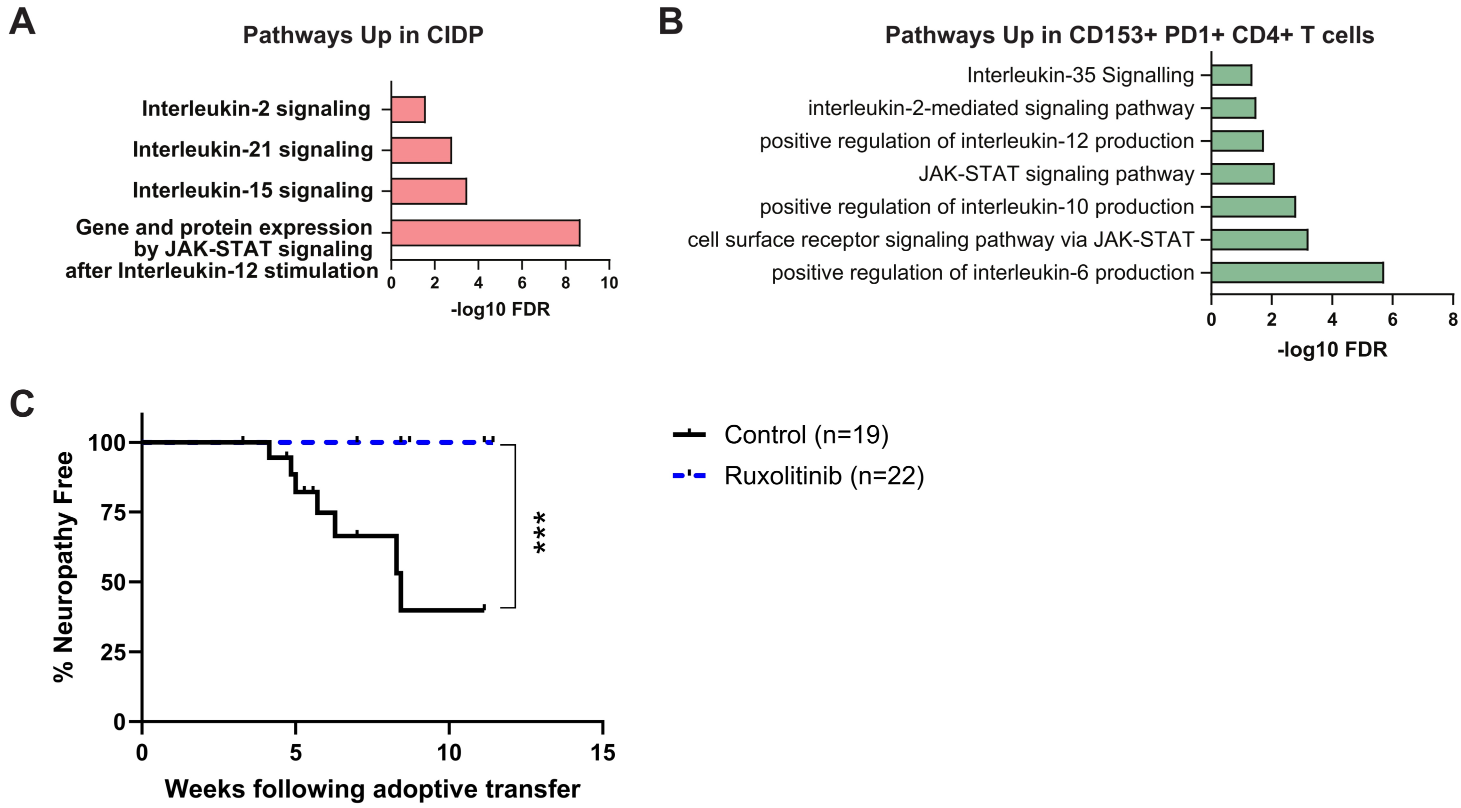


**Figure S5: JAK inhibitor Ruxolitinib is a senomorphic agent that protects against autoimmune peripheral neuropathy. A)** Gene Ontology analysis for upregulated genes in CD4+ T cells of CIDP patients in comparison to healthy donors. **B)** Gene Ontology analysis of upregulated genes in CD153+ PD1+ vs. CD153- PD1- CD4+ sorted T cells from *NOD.Aire*$^{GW/+}$ spleen. **C)** Kaplan-Meier curve of neuropathy incidence following adoptive transfer of stimulated splenocytes from *NOD.Aire*$^{GW/+}$ mice. Mice were randomly assigned to ruxolitinib or vehicle treatment (p=0.0003, Log-Rank test).
