## Supplemental Tabel 1-10 for "Age-associated Autoimmunity Driven by T cell Immunosenescence": Supp Table 8 .docx

| Antibody/Dye | Vendor | Fluorophore | Clone | Cat number |
| --- | --- | --- | --- | --- |
| Annexin V | Biolegend | FITC |  | 640906 |
| Fixable Viability Dye | Thermo Fisher Scientific | eFluor450 |  | 65-0863-14 |
| Human Antibodies | | | | |
| Anti human CD3 | Biolegend | APC/Cy7 | UCHT1 | 300426 |
| Anti human CD4 | Biolegend | PE | RPA-T4 | 300550 |
| Anti human CD8 | Biolegend | BV711 | SK1 | 344734 |
| Mouse Antibodies | | | | |
| Anti CD45 | Biolegend | AF700 | 30-F11 | 103128 |
| Anti p21 | abcam | PE | EPR18021 | ab314287 |
| Anti γH2AX | Biolegend | PECy7 | 2F3 | 613420 |
| Anti CD3 | Biolegend | AF700 | 17A2 | 100216 |
| Anti CD4 | Biolegend | Pacific Blue/FITC | GK1.5 | 100428  100406 |
| Anti CD4 | Biolegend | PECy7 | RM4-5 | 100528 |
| Anti CD4 | Invitrogen | APC | GK1.5 | 17-0041-82 |
| Anti CD4 | Biolegend | BV605 | RM4-5 | 100548 |
| Anti-CD153 | Thermo Fisher Scientific | PE | RM153 | 12-1531-82 |
| Anti-PD1 | Thermo Fisher Scientific | SuperBright 600 | J43 | 63-9985-82 |
| Anti-CD44 | Invitrogen | PECy7 | IM7 | 25-0441-82 |
| Anti-IFNγ | Biolegend | BV785 | XMG1.2 | 505837 |
| Anti-IL10 | BD Bioscience | FITC | JES5-16E3 | 554466 |
| Anti-IL21 | Invitrogen | PE | Mhalx21 | 12-7213-82 |
| Anti TNFα | Biolegend | PECy7 | MP6-XT22 | 506324 |
| Anti-p21 | Thermo Fisher Scientific | - | R.229.6 | MA5-14949 |
| Anti p75-NGFR | abcam | - | EP1039Y | ab52987 |
| Secondary antibodies | | | | |
| Anti rabbit IgG | abcam | AF647 | polyclonal | Ab150079 |
